# ΔNp63 Regulates Homeostasis, Stemness, and Suppression of Inflammation in the Adult Epidermis

**DOI:** 10.1101/2022.08.17.504172

**Authors:** Christopher E. Eyermann, Xi Chen, Ozge S. Somuncu, Jinyu Li, Alexander N. Joukov, Jiang Chen, Evguenia M. Alexandrova

## Abstract

The p63 transcription factor is critical for epidermis formation in embryonic development, but its role in the adult epidermis is poorly understood. Here we show that acute genetic ablation of ΔNp63, the main p63 isoform, in adult epidermis disrupts keratinocyte proliferation and self-maintenance and, unexpectedly, triggers an inflammatory psoriasis-like condition. Mechanistically, single-cell RNA sequencing revealed down-regulation of the cell cycle genes, up-regulation of differentiation markers, and induction of several pro-inflammatory pathways in ΔNp63-ablated keratinocytes. Intriguingly, ΔNp63-ablated cells disappear three weeks post-ablation, at the expense of the remaining non-ablated cells. This is not associated with active cell death mechanisms, but rather with reduced self-maintenance capacity. Indeed, *in vivo* wound healing assay, a physiological readout of the epidermal stem cell function, is severely impaired in ΔNp63-ablated mice. We found that the Wnt signaling pathway (Wnt10a, Fzd6, Fzd10) and the AP1 factors (JunB, Fos, FosB) are the likely ΔNp63 effectors responsible for keratinocyte proliferation/stemness and suppression of differentiation, respectively, while interleukins IL-1a, IL-18, IL-24, and IL-36γ are the likely negative effectors responsible for the suppression of inflammation. These data establish ΔNp63 as a critical node that coordinates epidermal homeostasis, stemness, and suppression of inflammation in the adult epidermis, upstream of known regulatory pathways.

## INTRODUCTION

Epidermis, the outermost skin layer, is responsible for the skin’s barrier function against a hostile environment and is composed of keratinocytes (KCs), the main cell type. One of the epidermal master regulators during embryonic development is the *p63* gene (Vanbokhoven et al. 2011), but its role in adult epidermis is poorly understood. p63 knockout (KO) embryos have severe defects of epidermis and its appendages, which is attributed to aberrant epidermal specification or KC self-maintenance, resulting in perinatal lethality (Mills et al. 1999; Yang et al. 1999). The *p63* gene produces multiple N- and C-terminal isoforms: full-length TAp63 and the N-terminally truncated ΔNp63 transcribed from an alternative internal promoter. ΔNp63 is the principal p63 isoform, evident by its significantly higher expression than TAp63 (Yang et al. 1999; Romano et al. 2009) and by the phenotype of ΔNp63-KO embryos that greatly mimic global p63-KOs and are also perinatally lethal (Romano et al. 2012; Chakravarti et al. 2014). p63/ΔNp63 continues to be strongly expressed in adult epidermis, and there in the basal/supra-basal gradient consistently with the epidermal stem cell (SC) niche (Yang et al. 1998; Parsa et al. 1999). Mutations in the *p63* gene are associated with familial EEC (ectrodactyly, ectodermal dysplasia, cleft lip/palate) and other skin fragility syndromes, highlighting a critical role of p63/ΔNp63 in the adult skin (Soares and Zhou 2018). However, perinatal lethality of KOs complicates *in vivo* adult studies, while knockdown (KD) reports yield conflicting results (Truong et al. 2006; Koster et al. 2007). Here, we investigated the role of ΔNp63 in adult epidermis using a robust, conditionally inducible mouse model and revealed its several critical roles there that we mechanistically linked to novel and previously identified ΔNp63 effector genes.

## RESULTS

### The Efficiency and Phenotype of ΔNp63 Ablation in the Adult Skin

To investigate the role of ΔNp63 in adult epidermis *in vivo*, we generated conditionally inducible ΔNp63^fl/fl^;K14-CreT2 mice, combining previously described ΔNp63^fl/fl^ (Chakravarti et al. 2014; Venkatanarayan et al. 2015; Napoli et al. 2022) and Krt14-CreERT2 alleles (Vasioukhin et al. 1999), and two control cohorts: ΔNp63^fl/fl^ (no Cre) and ΔNp63^fl/+^;K14-Cre (for heterozygous ablation) (Figure 1a). This allows for Tamoxifen-induced ΔNp63 ablation in the basal epidermal layer, its predominant expression site. Indeed, Tamoxifen significantly reduced *ΔNp63* and *total p63* mRNA levels in the skin of ΔNp63^fl/fl^;K14-CreT2 but not control mice, at 1 day and 1 wk post-ablation, which was restored at 3 wks (Figure 1b). Interestingly, *TAp63* was slightly upregulated, consistently with previous reports of ΔNp63 KD (Keyes et al. 2005; Truong et al. 2006; Venkatanarayan et al. 2015), but since is expression is minimal to begin with, its impact is probably minor. At the protein level, p63 (in lieu of ΔNp63) was greatly reduced in the interfollicular epidermis and less in the hair follicles (Figure 1c), likely due to their weaker Krt14 expression (Dai et al. 2011). Phenotypically, 77% of ΔNp63-ablated mice developed patchy, albeit temporary alopecia (Figure 1d, e; Supplementary Table 1), and many exhibited temporary loss of body weight (Supplementary Figure 1a, b).

**Figure 1.**
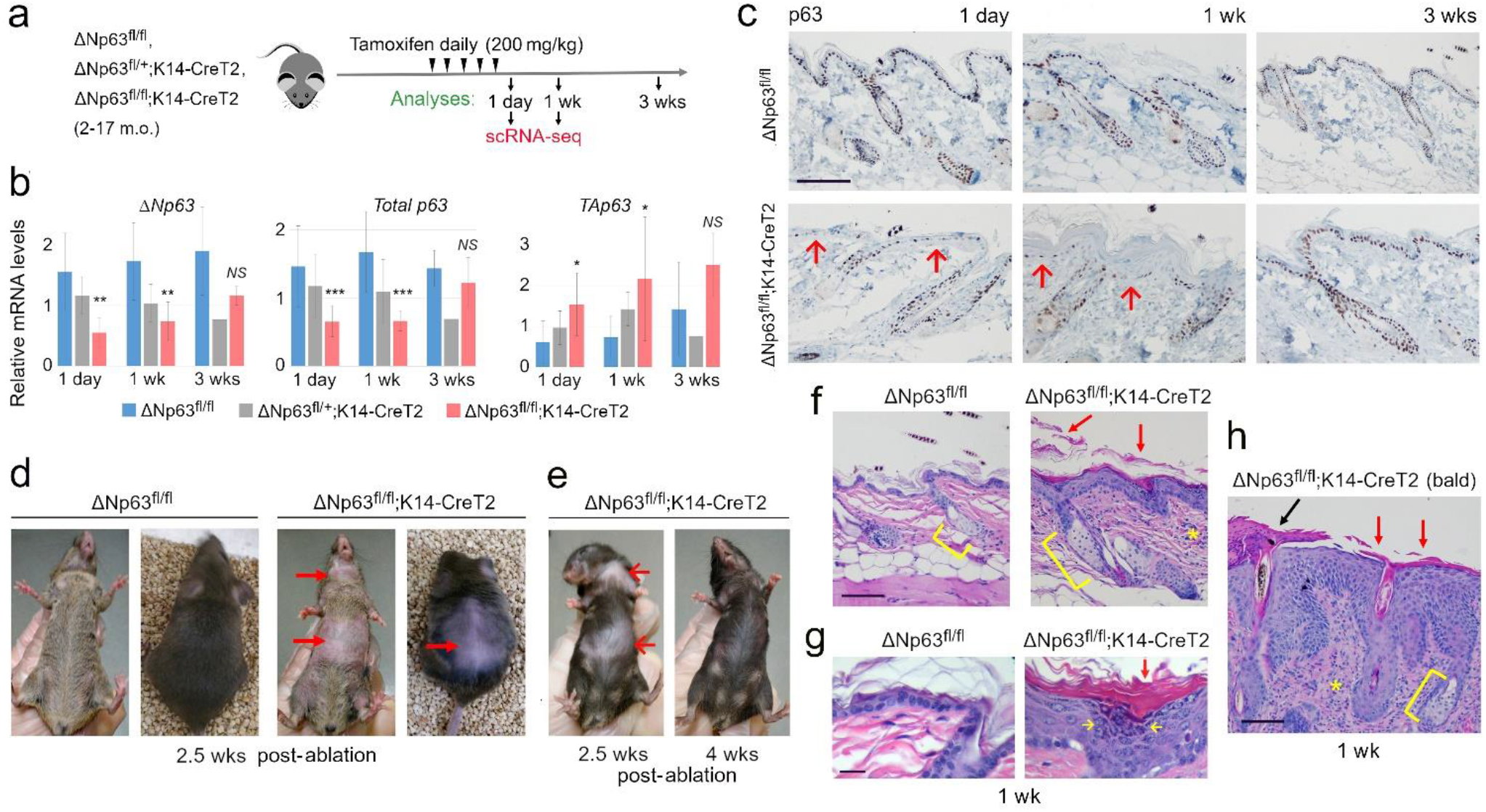
The efficiency and phenotype of ΔNp63 ablation in the adult skin. **a**. Experimental diagram. **b**. qRT-PCR for *ΔNp63*, total *p63* and *TAp63*, mean ± SD, *p<0.05, **p<0.01, ***p<0.001, *NS*, not significant (vs. ΔNp63^fl/fl^). **c**. IHC for p63 at the indicated time points, back skin. *Arrows*, interfollicular epidermis with absent p63. **d, e**. Temporary alopecia (*arrows*). **f-h**. H&E at 1 wk post-ablation showing acanthosis (epidermis thickening), hyperkeratosis (*red arrows*), enlarged keratohyalin granules (*yellow arrows*), parakeratosis (*black arrow*), fibrosis and immune cells infiltration (*asterisks*), and hyperplasia of the sebaceous gland (*brackets*). “Bald”, skin with alopecia. Scale bars, 100 µm (**c, f, h**), 25 µm (**g**).

Morphologically, there were no discernable defects at 1 day, however, numerous pathologies were noted at 1 wk: acanthosis (epidermal thickening), hyperkeratosis (thickened cornified layer indicative of excessive keratin production and shedding), enlarged keratohyalin granules, parakeratosis (aberrant shedding of nucleated cells), and changes in the dermis - fibrosis, inflammatory infiltration, and hyperplasia of the sebaceous glands (Figure 1f-h; Supplementary Figure 1c-f), resembling psoriasis vulgaris in humans, except for hyperplastic sebaceous glands (hypoplastic in psoriasis) (Rittié et al. 2016). While alopecia may or may not accompany psoriasis in humans (George et al. 2015; Rittié et al. 2016), since alopecic (“bald”) skin strongly correlated with abnormal histopathology in ΔNp63-ablated mice (Figure 1h; Supplementary Figure 1e), we speculate it is not a separate phenotype. Since ΔNp63 mRNA and protein expression is restored at 3 wks post-ablation and so are histologic abnormalities, followed by the reversal of alopecia at 4 wks (Figure 1e), this suggests a rapid loss of ΔNp63-ablated KCs at the expense of the remaining non-ablated KCs and their progeny. We speculate that if the ablation efficiency was 100%, this would be incompatible with the existence of epidermis, similarly to p63/ΔNp63-KO embryos.

### Proliferation and Self-Maintenance Defects of ΔNp63-Ablated Keratinocytes

Epidermal thickening (acanthosis) in ΔNp63-ablated mice suggested increased KC proliferation, which we confirmed by increased Ki67 positivity at 1 wk, but not at 1 day or 3 wks post-ablation (Figure 2a, b; Supplementary Figure 1g, h). However, serial sections revealed that hyper-proliferation was mostly confined to the remaining non-ablated (ΔNp63^pos^) KCs, while ΔNp63^neg^ KCs were hypo-proliferative (Figure 2c, d). In agreement, the alopecic skin patches were always both, Ki67^pos^ and p63^pos^ (Figure 2e). This suggests that ΔNp63 ablation non-cell-autonomously induces proliferation in the remaining ΔNp63^pos^ KCs, e.g., via secreted pro-inflammatory cytokines (see below).

**Figure 2.**
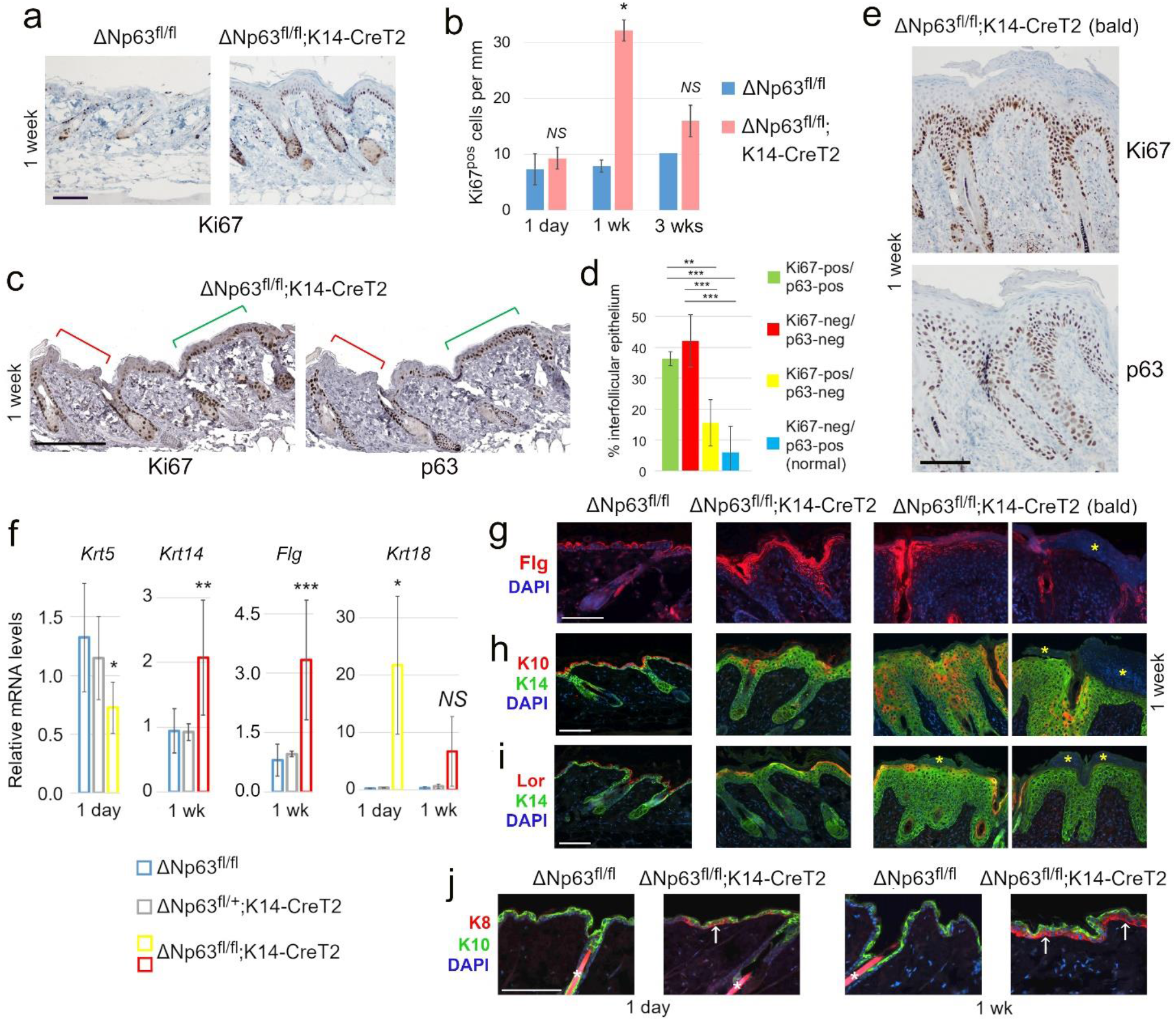
Proliferation and differentiation defects of ΔNp63-ablated skin. **a, b**. Ki67 IHC (**a**) and quantification (**b**). **c, d**. Ki67 and p63 IHC on serial sections (**c**) and quantification from n=5 mice (**d**). *Red bracket*, Ki67^neg^/p63^neg^ area; *green bracket*, Ki67^pos^/p63^pos^ area. **e**. Ki67 and p63 IHC in alopecic skin. **f**. qRT-PCR of select stratification markers, mean ± SD vs. ΔNp63^fl/fl^. **g-j**. Immunofluorescent staining at 1 wk post-ablation (except **j**). *Yellow asterisks*, parakeratosis corresponding to the areas of lost Flg, K10 and Lor expression; *arrows*, aberrant Krt8 upregulation; *white asterisks*, non-specific hair shaft staining (**j**). “Bald”, skin with alopecia. Scale bars, 100 µm. (**b, d, f**) Mean ± SD, *p<0.05, **p<0.01, ***p<0.001, *NS*, not significant.

Reduced proliferation and rapid loss of ΔNp63^neg^ KCs within 3 wks post-ablation suggests that they either die (if ΔNp63 is required for KC survival) or have reduced self-maintenance and disappear via losing competition to the remaining - better fit - ΔNp63^pos^ KCs. We found no evidence of active cell death: necrosis (RIP3) and apoptosis (TUNEL, Cleaved Caspase-3) markers were not significantly increased and, notably, localized to the outermost layer, not the basal ΔNp63-ablated KCs (Supplementary Figure 2 and *not shown*), a cell death pattern characteristic for psoriatic KCs (Iizuka et al. 2004). Likewise, the markers of cellular senescence (SA-X-gal, p16) were not induced (*not shown*). Altogether, these data indicate that ΔNp63 is required for KC self-maintenance and optimal fitness rather than survival.

### Differentiation and Stratification Defects of ΔNp63-Ablated Epidermis

Hyperkeratosis and increased keratohyalin production (Figure 1f-h) suggest accelerated terminal differentiation. Indeed, analysis of stratification markers revealed upregulation of the granular marker Flg and downregulation of the basal marker Krt5, a ΔNp63 target gene (Romano et al. 2009), whereas Lor and the spinous marker Krt10 were not affected (Figure 2f-i; Supplementary Figure 3a-c, h, and *not shown*). In contrast, basal Krt14, also a reported ΔNp63 target (Romano et al. 2007; Romano et al. 2009), was not reduced and rather up-regulated at 1 wk post-ablation, expanding into the supra-basal layers and overlapping with the spinous and granular markers Krt10 and Lor, respectively (Figure 2f, h, i). Furthermore, ΔNp63-depleted KCs were previously shown to spontaneously reprogram into induced pluripotent stem cells (iPSC) via a micro-RNA processing factor DGCR8 (Chakravarti et al. 2014). We found no induction of iPSC markers, but a significant de-repression of immature epidermal markers Krt8/18, as previously reported (Truong et al. 2006; Shalom-Feuerstein et al. 2011; Romano et al. 2012; Chakravarti et al. 2014; Fan et al. 2018), and only minor down-regulation of *DGCR8* at the late 1 wk time-point (Figure 2f, j; Supplementary Figure 3d-g), suggesting that DGCR8 is not a major ΔNp63 effector in adult epidermis. Altogether, these data indicate a critical role of ΔNp63 in stratification of adult epidermis, suppression of KC differentiation, and repression of embryonic keratins.

### Molecular Pathways Altered in ΔNp63-Ablated Epidermis

To gain insight into the molecular mechanisms of the observed defects and zoom into ΔNp63^neg^ vs. the remaining ΔNp63^pos^ KCs, we performed single-cell RNA sequencing (scRNA-seq) analysis of ΔNp63-ablated vs. control KCs at 1 day and 1 wk post-ablation (from n=3 pooled mice each). At 1 day, ΔNp63-ablated KCs segregated into two distinct clusters: (1) with a gene expression signature similar to control KCs (average *p63* mRNA expression=0.36 of control) and (2) with a completely new gene expression signature (*p63* mRNA=0.18 of control), which we termed p63^med^ and p63^low^ clusters, respectively (Figure 3a; Supplementary Figure 4a, b). At 1 wk, only a minority of KCs had a gene expression signature similar to control KCs (C1-C3), while the majority had a distinct gene expression signature and elevated *Ki67* expression, therefore, we termed them “Normal-like” and “Hyper-proliferative”, respectively (Figure 3b; Supplementary Figure 4a, b). Notably, *p63* expression likely reflects *ΔNp63* (not directly detectible by scRNA-seq), since known ΔNp63 target genes were down-regulated, while TAp63 target genes were upregulated in ΔNp63-depleted cells (Figure 3c; Supplementary Figure 3c).

**Figure 3.**
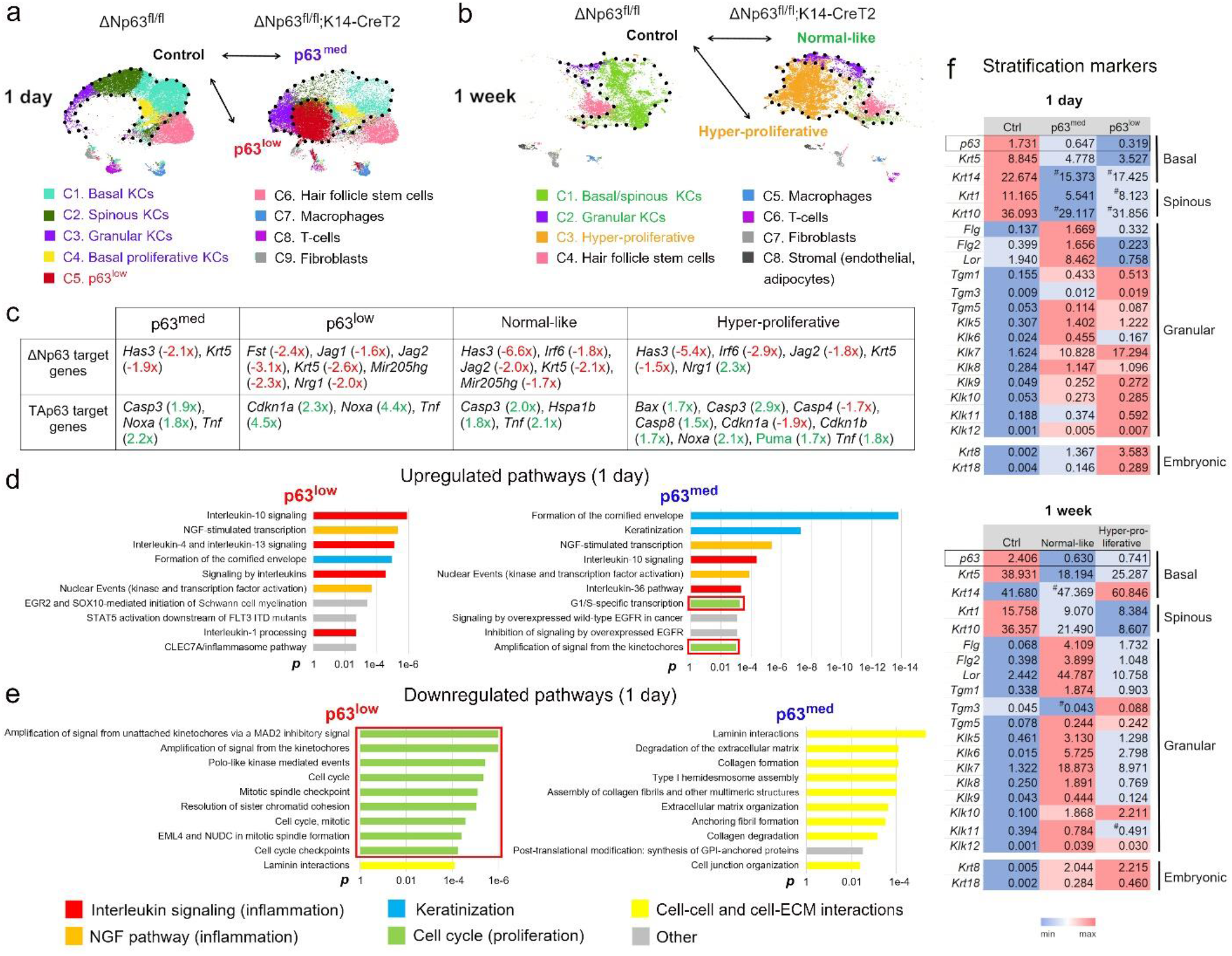
ScRNA-seq of ΔNp63-ablated KCs reveals altered molecular pathways. **a, b**. UMAP plots. At 1 day post-ablation, clusters C1-C4 (collectively, p63^med^) and C5 (p63^low^) were separately compared with control clusters C1-C4 to reveal differentially expressed genes and pathways. At 1 wk post-ablation, clusters C1-C2 (collectively, “Normal-like”) and C3 (“Hyper-proliferative”) were separately compared with control clusters C1-C2. **c**. Relative expression levels ΔNp63 and TAp63 target genes in the indicated clusters. **d, e**. The top ten upregulated (**d**) and down-regulated (**e**) pathways at 1 day post-ablation, Reactome. **f**. Heatmaps of stratification markers at 1 day and 1 wk post-ablation. ^#^, not significant. Note, at 1 wk, ΔNp63-ablated KC likely reside in the “Normal-like” cluster, reflected by their gene expression signature similar to ΔNp63^low^ at 1 day.

Analysis of differentially expressed genes and the Reactome pathways at 1 day revealed that the most significantly upregulated pathways in ΔNp63-ablated KCs are (1) “Keratinization” (including differentiation genes), (2) Interleukin signaling, and (3) the Nerve Growth Factor (NGF) pathway (Figure 3d), the latter two linked to inflammatory skin conditions, see below. The most down-regulated pathways shared by ΔNp63^low^ and ΔNp63^med^ clusters were the cell-cell and cell-ECM (extracellular matrix) interaction pathways (Figure 3e), consistently with previous reports (Guan et al. 2021). Notably, ΔNp63^low^ and ΔNp63^med^ clusters showed an opposite regulation of the cell cycle genes and previously identified “core” stemness genes (Wang et al. 2020): down-regulation in ΔNp63^low^ and upregulation in ΔNp63^med^ (Figure 3d, e, *red boxes*; Supplementary Figure 5a). This mechanistically explains why ΔNp63-ablated KCs are hypo-proliferative with reduced self-maintenance, while ΔNp63^pos^ KCs (likely in the p63^med^ cluster) are hyper-proliferative. ScRNA-seq analysis at 1 wk revealed similar pathway changes, except for a widespread upregulation of the cell cycle genes exclusively in the “Hyper-proliferative” cluster (Supplementary Figure 5b, c), suggesting that it represents the expanded “psoriatic”/ΔNp63^pos^ KCs, whereas ΔNp63^low^ KCs likely belong to the “Normal-like” cluster.

Notably, 1 day and 1 wk scRNA-seq revealed expansion of the granular cluster (Figure 3a, b, *purple*) and a shift from the basal to the granular layer identity evident by upregulation of the differentiation genes *Flg, Tgm1/3/5, Klk5-12* (Figure 3f) as part of the “Keratinization” pathways. This independently confirmed accelerated terminal differentiation upon ΔNp63 ablation. Overall, scRNA-seq in an unbiased way revealed four major molecular defects in ΔNp63-ablated adult epidermis: impaired proliferation and stemness; enhanced terminal differentiation (together, imbalanced homeostasis); upregulation of the inflammatory pathways; and reduced cell-cell and cell-ECM interactions.

### ΔNp63 Effectors Responsible for Epidermal Homeostasis

Many pathways are known to regulate epidermal homeostasis, e.g., the Wnt/β-catenin and Shh pathways responsible for KC proliferation, stemness and maintenance (Clevers et al. 2014; Abe and Tanaka 2017) and the Notch pathway and AP1 transcription factors (aka Jun/Fos) responsible for KC differentiation (Eckert et al. 2013; Nowell and Radtke 2013). Analysis of these and other signaling pathways revealed significant down-regulation of the Wnt pathway, as well as Shh and Edaradd components, in the most depleted ΔNp63^low^ and Normal-like clusters (Figure 4a, d; Supplementary Figure 6a). In addition, *Col17a1* from the cell-ECM interaction pathways previously shown to regulate KC proliferation and stemness and a reported ΔNp63 target gene in squamous cell carcinoma (Ramsey et al. 2013; Matsumura et al. 2016; Watanabe et al. 2017; Liu et al. 2019), was strongly down-regulated, >4 folds (Figure 4b; Supplementary Figure 6b). On the contrary, several AP1 transcription factors were strongly upregulated (Figure 4c, e). The other signaling pathways, e.g., Notch, FGF, TGFβ, including previously proposed to mediate p63/ΔNp63 functions (Ramsey et al. 2013; Fan et al. 2018), were not consistently affected (Supplementary Figure 6a). Analysis of a previously published p63 ChIP-seq database (Kouwenhoven et al. 2015) revealed potential p63 binding sites in several Wnt pathway components (*Fzd6, Fzd10, Wnt10a*), *Col17a1*, and several AP1 genes (*JunB, Fos, FosB, Fosl2/Fra1*, Figure 4f, g; Supplementary Figure 6c). We confirmed direct binding of p63 to many of these sites by ChIP-qPCR (Figure 4h). Since ΔNp63 can both, activate and repress its target genes (Kouwenhoven et al. 2015; Sethi et al. 2017; Fan et al. 2018; Napoli et al. 2022), these data strongly suggest that ΔNp63 coordinates epidermal homeostasis by activating the pro-proliferative/pro-stemness Wnt pathway and *Col17a1*, while repressing the differentiation driving AP1 factors.

**Figure 4.**
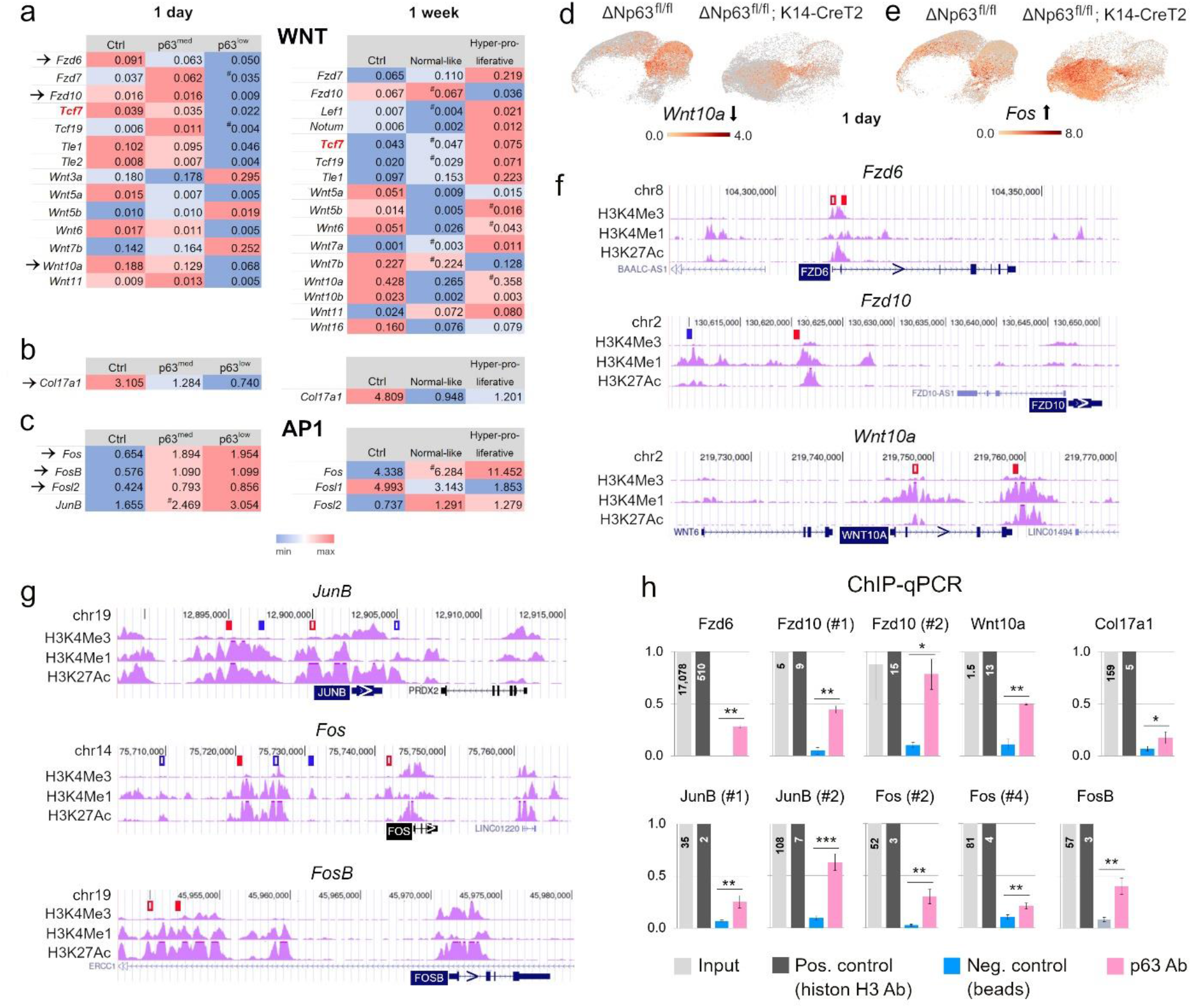
ΔNp63 effectors responsible for epidermal homeostasis. **a-c**. The heatmaps of the differentially expressed genes from the pro-proliferative/pro-stemness Wnt pathway (**a**) and *Col17a1* (**b**) and pro-differentiation AP1 genes (**c**). *Tcf7* (*red*) is a Wnt target gene and serves as the pathway readout. ^#^, not significant. **d, e**. Representative UMAP plots. **f, g**. Potential p63 binding sites in the Wnt pathway components (**f**) and AP1 genes (**g**) from a published p63 ChIP-seq database overlayed with chromatin modification tracks (*purple*) in the UCSC genome browser, both in NHEK cells. *Red*, in all conditions; *blue*, only in proliferative conditions; *solid*, sites confirmed by ChIP-qPCR; not to scale. **h**. ChIP-qPCR of p63 binding in HaCaT cells, *p<0.05, **p<0.01, ***p<0.001.

### Impaired Wound Healing in ΔNp63-Ablated Skin

One physiologically important role of epidermal SCs is to enable wound healing (Yang et al. 2019). Therefore, we tested whether reduced stemness/self-maintenance of ΔNp63-ablated KCs would affect their wound healing capacity. We found that, indeed, full-skin excision wounds introduced at 1 day post-ablation did not heal in ΔNp63-ablated mice (Figure 5a, b), failed to form the hyper-thickened epidermal ridge despite acanthosis away from the wound (Figure 5c, d) and to upregulate Col17a1 (Figure 5e). Notably, proliferation was not affected near the wound (Figure 5f, g), despite an increase away from the wound as aforementioned, and even non-ablated p63^pos^ cells failed to migrate towards the wound (Figure 5h) as one might expect from other incomplete KOs (Yan et al. 2021). This reveals a dominant-negative effect of ΔNp63^neg^ KCs towards ΔNp63^pos^ KCs, which we speculate is due to defective cell-ECM interactions. Overall, this reveals a critical, physiologically relevant role of ΔNp63 as a SC regulator in adult epidermis.

**Figure 5.**
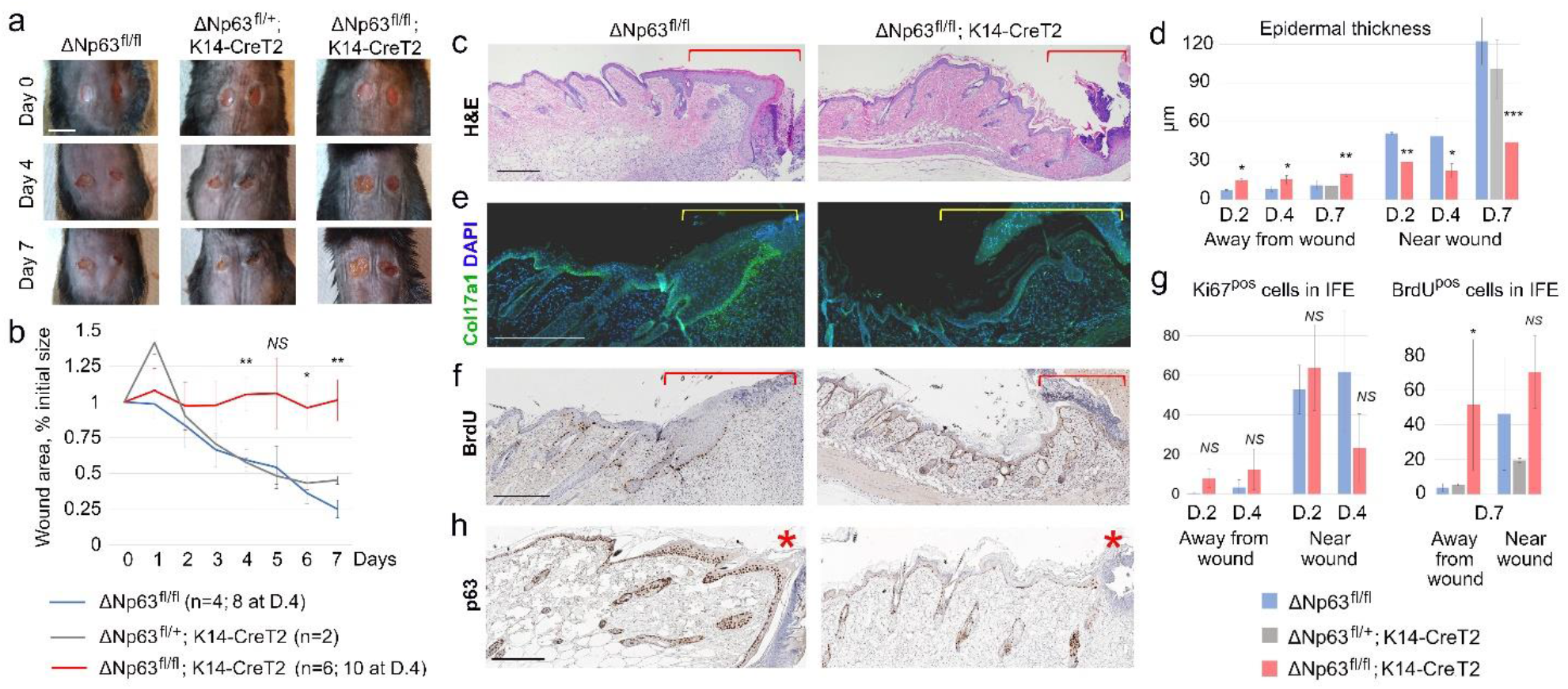
Defective wound healing in ΔNp63-ablated skin. **a, b**. Representative images (**a**) and quantification of the wound area over time (**b**). **c, d**. Impaired formation of the hyper-thickened epidermal ridge, H&E (**c**) and quantification (**d**). **e**. Immunofluorescent staining for Col17a1. **f, g**. IHC for BrdU (**f**) and quantification of Ki67^pos^ (*not shown*) and BrdU^pos^ cells per five interfollicular epidermis (IFE) spaces (**g**). **h**. IHC for p63. All images are at day 7, except **h** (day 4). *Brackets, asterisks*, the wound area. (**b, d, g**) Mean ± SD, \**p*<0.05, \*\**p*<0.01, \*\*\**p*<0.001, *NS*, not significant (vs. ΔNp63^fl/fl^). Scale bars, 1 cm (**a**), 200 µm (**c, e, f, h**).

### Regulation of Inflammation by ΔNp63

Consistently with the inflammatory phenotype (Figure 1f, h), scRNA-seq revealed upregulation of several interleukin pathways: IL-1, IL-4/13, IL-10, IL-36, and the NGF pathway (Figure 3d) associated with inflammatory skin conditions such as psoriasis and AD (Pincelli 2000; Guttman-Yassky et al. 2011; Johnston et al. 2017) (Figure 6a, b; Supplementary Figure 7a). In addition, many epidermis-specific “alarmin” genes were upregulated: *Krt6a/b, Krt16/17, Lce3c-f, s100a8/9, Slurp1/2, Sprr1/2* - also associated with psoriasis and occasionally AD (Figure 6c). Several lines of evidence point to a psoriasis-like rather than AD-like condition in ΔNp63-ablated mice: upregulation of *IL-1, IL-36* and *s100a8/9* (higher in psoriasis than AD (Kerkhoff et al. 2012; D’Erme et al. 2015)) and “hallmark of psoriasis” *Krt6/16/17* genes (Zhang et al. 2019), increased differentiation/cornification (reduced in AD), characteristic parakeratosis, involvement of Tnf and Ifngr1/2 (Supplementary Figure 7a) (Trzeciak et al. 2017; Albanesi et al. 2018). Moreover, analysis of human plaque psoriasis biopsies revealed protein expression patterns similar to those in ΔNp63-ablated mouse skin: IL-36 upregulation in the outermost layer, Krt14 expansion into the supra-basal layers, and reduced Flg expression (Iizuka et al. 2004) (Figure 6d-h, compare to Figure 2g-i). This suggests that ΔNp63 is an epidermis-specific negative regulator of psoriatic inflammation. Analysis a p63 ChIP-seq database (Kouwenhoven et al. 2015) revealed potential p63 binding sites in several cytokines from the upregulated inflammatory pathways: *IL-1a*, a previously established ΔNp63 target (Barton et al. 2010), *IL-18, IL-24, IL-36γ*, and *Ifng1/2* (Figure 6i and *not shown*) - all implicated in psoriasis (Mee et al. 2006; Rasmy et al. 2011; Wang et al. 2017; Albanesi et al. 2018; Mitamura et al. 2020). We confirmed direct binding of p63 to many of these sites by ChIP-qPCR (Figure 6j). Finally, we found a significantly decreased number of p63^pos^ KCs in human psoriatic biopsies compared to normal and non-lesional skin (Figure 6k, l), consistently with previous reports (Gu et al. 2006; Li et al. 2014). Altogether, this establishes ΔNp63 as a critical suppressor of psoriatic inflammation in the adult skin, likely by directly repressing select cytokine genes, a completely novel role not inferred from previous studies.

**Figure 6.**
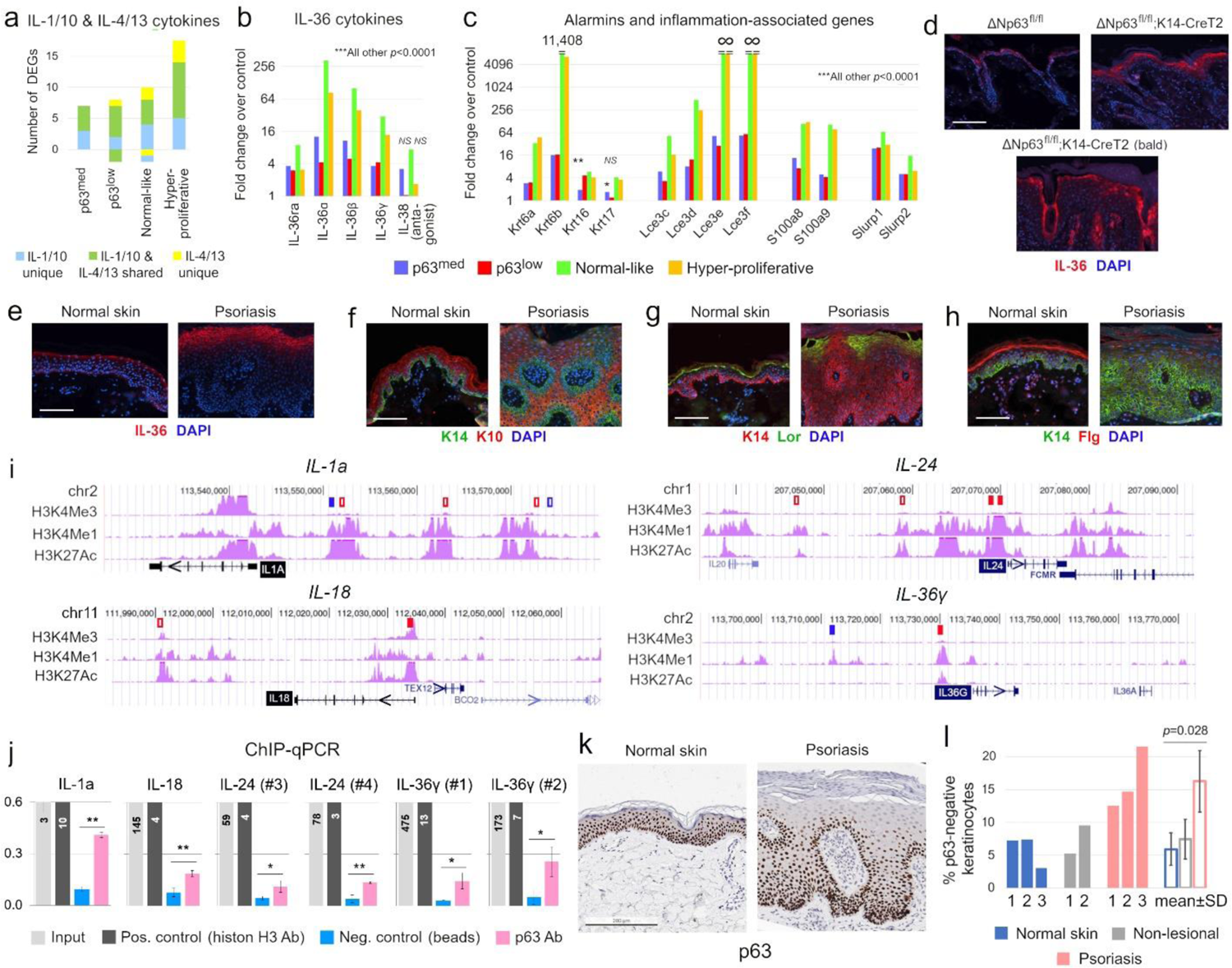
ΔNp63 as the suppressor of psoriatic inflammation. **a**. The total number of significantly altered *IL-1/10* and *IL-4/13* genes. **b, c**. Upregulation of *IL-36* genes (**b**) and epidermal alarmins (**c**). “∞”, values were divided by zero. **d-h**. Immunofluorescence in ΔNp63-ablated mouse skin (**d**) and human psoriatic biopsies (**e-h**). **i**. Potential p63 binding sites in the interleukin genes from a p63 ChIP-seq database overlayed with chromatin modifications (*purple*), NHEK cells; not to scale. *Red*, all conditions; *blue*, proliferative conditions; *solid*, confirmed by ChIP-qPCR. **j**. ChIP-qPCR of p63 binding in HaCaT cells. **k, l**. p63 IHC in human psoriatic biopsies (**k**) and quantification of p63^neg^ KCs (**l**). Scale bars, 100 µm (**d-h**), 200 µm (**l**). (**b, c, j**) \**p*<0.05, \*\**p*<0.01, *NS*, not significant; all others, *p*<0.001 (**b, c**).

## DISCUSSION

Here we uncovered that ΔNp63 plays several critical roles in adult epidermis: (1) promotes proliferation, self-maintenance and stemness of basal KCs, thus contributing to wound healing, (2) suppresses KC differentiation, and (3) suppresses epidermal inflammation (Supplementary Figure 8a), all of which are perturbed in ΔNp63-ablated epidermis (Supplementary Figure 8b). While it is still debatable whether p63/ΔNp63 regulates epidermal development via KC stemness/self-maintenance (Yang et al. 1999; Pellegrini et al. 2001; Senoo et al. 2007; Romano et al. 2012) or commitment/differentiation (Mills et al. 1999; Koster et al. 2004; Truong et al. 2006; Koster et al. 2007; Shalom-Feuerstein et al. 2011), our data strongly support the former, at least in the adult skin. Similarly, there is an ongoing debate whether p63/ΔNp63 promotes (Parsa et al. 1999; Pellegrini et al. 2001; Truong et al. 2006; Senoo et al. 2007; Romano et al. 2012) or represses KC proliferation (Koster et al. 2007; Chakravarti et al. 2014), including in the adult models. Our data reveal that ΔNp63-depleted KCs are hypo-proliferative, while the remaining non-ablated KCs are hyper-proliferative, likely in response to paracrine inflammatory signals from the depleted KCs. This suggests that the previous reports of KC hyper-proliferation upon p63/ΔNp63 depletion also deal with incomplete KD. Regarding another intriguing phenotype of ΔNp63-ablated mice, loss of body weight, we excluded defects in the esophagus and tongue that might cause malnutrition (in fact, p63 was not down-regulated there; *not shown*) and speculate it is caused by salivary gland defects (Min et al. 2020).

Several signaling pathways downstream of ΔNp63 were found in embryonic and adult KCs: Wnt, Notch, FGF and Eda, while AP2-AP1 was proposed ΔNp63-independent (Ramsey et al. 2013; Sethi et al. 2017; Fan et al. 2018). Our data strongly support the role of the Wnt pathway (and possibly, Shh and Eda), but not others, as well as AP1 as ΔNp63 effectors. While we found the same Wnt receptors as previously reported (*Fzd6, Fzd10*), the ligand is different, *Wnt10a* as opposed to *Wnt3a, Wnt4, Wnt7b, Wnt10b* in the embryos (Fan et al. 2018). In addition, Col17a1, a transmembrane hemidesmosomal protein localized to the dermal surface of basal KCs, is likely a ΔNp63 effector that mediates both, epidermal SCs and cell-ECM interactions (Matsumura et al. 2016; Liu et al. 2019).

One novel function of ΔNp63 we uncovered is suppression of inflammation. Previously, ΔNp63 overexpression was shown to induce AD in mice (Du et al. 2014; Rizzo et al. 2016; Jiménez-Andrade et al. 2022) and to promote breast cancer via CXCL2 and CCL22 chemokines (Kumar et al. 2018). In contrast, our data revealed that ΔNp63 suppresses - not promotes - psoriasis-like epidermal inflammation. Psoriasis is a chronic autoimmune disease driven by an interplay between the immune system (Th1, Th17 cells and their secreted cytokines) and epidermis, and the primary role of KCs over the immune cells has been recently proposed (Albanesi et al. 2018; Ali et al. 2020; Zhou et al. 2022). Our findings support this unconventional model, evident by a quick and broad upregulation of inflammatory cytokines in ΔNp63-ablated KCs and reduced p63 expression in human psoriatic biopsies, consistently with previous reports of reduced expression of ΔNp63 mRNA, protein, and target genes in psoriasis and association of the *p63* genomic locus with susceptibility to psoriasis (Gu et al. 2006; Li et al. 2014; Yin et al. 2015). In sum, here we established ΔNp63 as the gatekeeper of adult epidermis via several critical functions: homeostasis, stemness, and suppression of inflammation.

## MATERIALS AND METHODS

### Mouse Strains, Isolation of Primary Keratinocytes

ΔNp63^fl/fl^ (Chakravarti et al. 2014) and Krt14-CreT2 mice (Jackson Labs, Bar Harbor, ME, strain 005107) were previously described (Vasioukhin et al. 1999). They were crossed to obtain experimental cohorts on a mixed 129SVJ/C57Bl6J/FVBN background. For ΔNp63 ablation, 2-17 m.o. mice received five doses of 200 mg/kg Tamoxifen (Sigma, Louis, MO, Cat. No. T5648-5G) in corn oil (Sigma, Cat. No. C8267) via daily oral gavage. All animals were treated humanely and according to the guidelines by the Stony Brook University Institutional Animal Care and Use Committee. Males and females were used at 1:1 ratio, no differences were noted between different genders or ages. Primary mouse KCs were isolated as previously described (Jensen et al. 2010), see “Supplementary Material” for details.

### Quantitative RT-PCR, Immunohistochemistry, Immunofluorescent Staining

Total RNA was extracted from whole back skin or from isolated KCs and qRT-PCR was performed as previously described (Eyermann et al. 2021). qPCR primers for total *p63*, Δ*Np63, TAp63* (Wolff et al. 2009), *HPRT, Oct4, Sox2, Nanog* (Nemajerova et al. 2012) were previously described. See “Supplementary Material” for additional primers. Immunohistochemistry (IHC) and immunofluorescent staining were previously described (Eyermann et al. 2021), see “Supplementary Material” for details and the primary antibodies.

### Single-Cell RNA Sequencing, Pathway Analysis, Chromatin Immunoprecipitation

ScRNA-seq was performed at NYU Genome Technology Center as previously described (Eyermann et al. 2021), except that Chromium Next GEM Single Cell 5′ Library & Gel Bead Kit v1.1, PN-1000165 was used at 1 wk post-ablation, and K-mean clustering (*k* = 10) was used for dimensionality reduction at both time-points. Pathway analysis was done using Reactome (www.reactome.com) based on the differentially expressed genes from scRNA-seq. For heatmaps, the genes with absolute expression levels <0.01 across the three assessed categories were excluded. Analysis of human genomic sequences and epigenetic regulatory tracks was done in UCSC Genome Browser, version GRCh37/hg19 (www.genome.ucsc.edu), using the ENCODE function to detect H3K4Me3 (promoters), H3K4Me1 (inactive enhancers) and H3K27Ac (active enhancers) in NHEK cells. These were overlayed with p63 binding sites at Day 0 (proliferation) and Day 7 (differentiation) in NHEK cells (Kouwenhoven et al. 2015). Chromatin Immunoprecipitation (ChIP) was performed on HaCaT cells using Ab500 kit (Abcam, Cambridge, UK) with p63 antibody Ab53039 (Abcam) and Histon H3 antibody Ab1791 (Abcam) as a positive control. See “Supplementary Material” for qPCR primers.

### Wound Healing Assay

For wound healing assay, mice were anesthetized with Ketamine/Xylazine (60 mg/kg; 100 mg/kg, respectively, in 0.67% NaCl), their back skin was shaved and disinfected, and two bilateral full-skin excision wounds were introduced with a sterile 6 mm biopsy punch. BrdU (Life Technologies, Carlsbad, CA, Cat. No. 000103) was i.p. injected two hrs prior to euthanasia. Epidermal thickness “Near wound” was measured at the thickest site excluding hair follicles. BrdU^pos^ and Ki67^pos^ cells “Near wound” and “Away from wound” were counted in the first five interfollicular spaces next to the wound and in interfollicular spaces 7 to 12, respectively. Two sections on both wound sides were analyzed (n=4 measurements per wound).

### Statistical Analysis

Unpaired two-tailed Student’s *t*-test was used in all experiments, except scRNA-seq and Reactome analyses. All experiments were performed in at least three biological replicas, unless indicated otherwise.

## DATA AVAILABILITY

ScRNA-seq data are available on Gene Expression Omnibus, accession number GSE213528.

## ACKNOWLEDGEMENTS

We thank Stony Brook Cancer Center’s Histology core facility for assistance with tissue histology, the Biobank for providing human skin biopsies, and the Biostatistics and Bioinformatics core facility for assistance with scRNA-seq analysis. We thank Dr. Elsa Flores for ΔNp63^fl/fl^ mice, Dr. Ute Moll and Dr. Daniel Lozeau for assistance with skin histopathology, Hemma Kilawan for assistance with data analysis, NYU Genome Technology Center for performing scRNA-seq experiments, and Histowiz for assistance with IHC staining. EMA was supported by NIH grant K22CA190653 and Stony Brook Cancer Center startup funds. JC was supported by NIH grant R01AR061485.

## AUTHOR CONTRIBUTIONS

Conceptualization: EMA (lead), JC (supporting). Investigation: CEE, XC, OSS, EMA. Formal analysis: JL, ANJ, EMA. Visualization: EMA (lead), ANJ (supporting). Writing - original draft preparation: EMA. Writing - review and editing: CEE, XC, OSS, JL, ANJ, JC, EMA.

