## Supplement for "ΔNp63 Regulates Homeostasis, Stemness, and Suppression of Inflammation in the Adult Epidermis"

### SUPPLEMENTARY FIGURES

**Supplementary Table 1**. Frequency of alopecia in ΔNp63-ablated and control adult mice.

| | $\Delta Np63^{fl/fl}$ | $\Delta Np63^{fl/+}$;K14-CreT2 | $\Delta Np63^{fl/fl}$;K14-CreT2 |
|---|---|---|---|
| Exp. 1 (4.5 m.o.) | 0/8 (0%) | *NA* | 5/7 (71%) |
| Exp. 2 (4.5 m.o.) | 0/6 (0%) | *NA* | 14/14 (100%) |
| Exp. 3 (9 m.o.) | 0/2 (0%) | *NA* | 1/4 (25%) |
| Exp. 4 (17 m.o.) | 0/4 (0%) | *NA* | 2/4 (50%) |
| Exp. 5 (2-5 m.o.) | *NA* | 0/1 (0%) | 2/3 (67%) |
| Exp. 6 (8-9 m.o.) | 0/4 (0%) | 0/4 (0%) | 3/3 (100%) |
| Total | 0/24 (0%) | 0/5 (0%) | 27/35 (77%) |

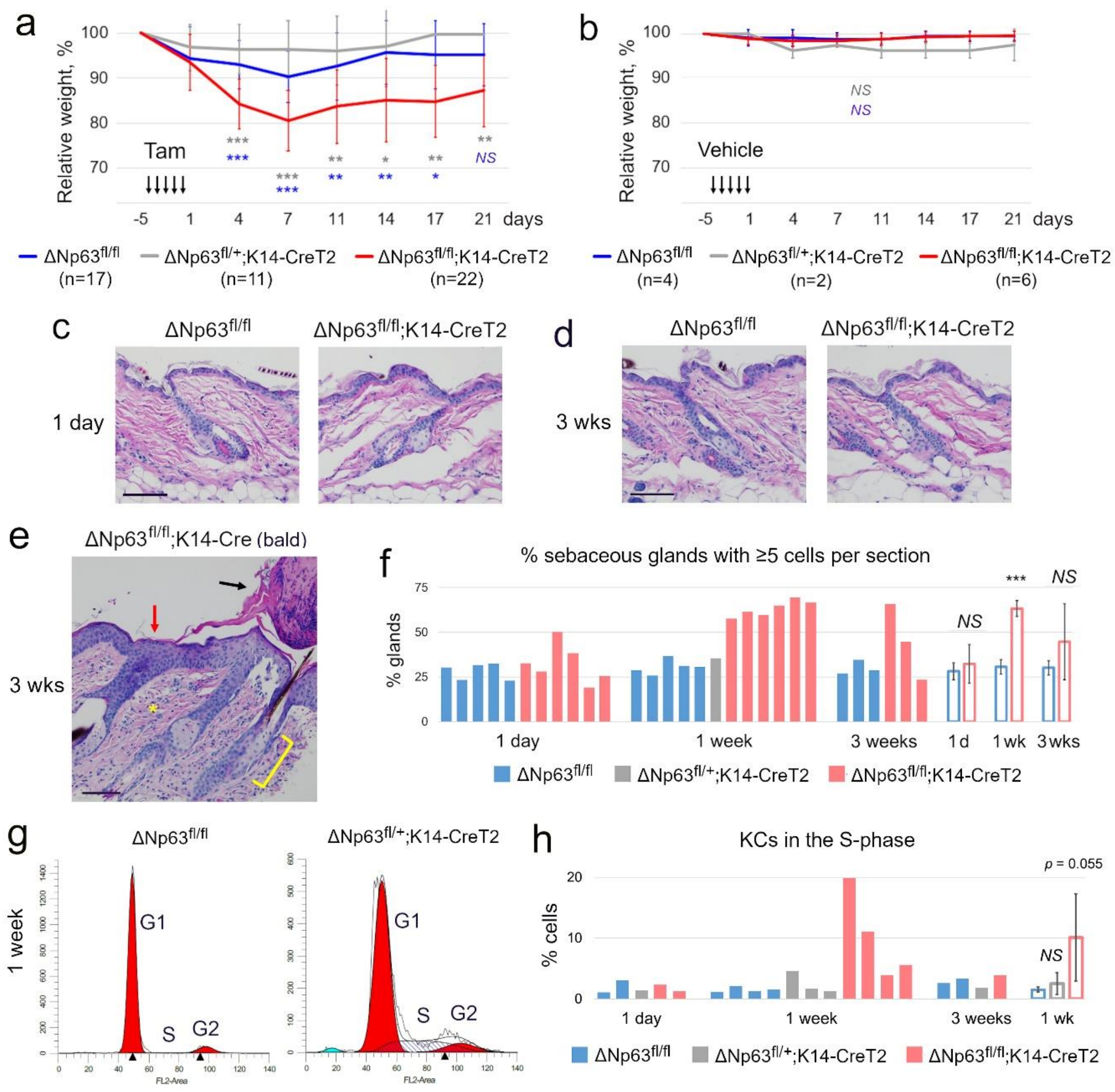


**Supplementary Figure 1**. Additional phenotypes of ΔNp63-ablated mice. **a, b.** Loss of body weight after Tamoxifen (**a**), but not Vehicle treatment (**b**); initial body weight was not different between the three phenotype (data not shown). **c-e.** H&E at 1 day (**c**) and 3 wks (**d, e**) post-ablation, including a rare remaining alopecic (“bald”) spot showing acanthosis, hyperkeratosis (*red arrow*), parakeratosis (*black arrow*), fibrosis and lymphocytic infiltration (*asterisk*), and hyperplasia of the sebaceous gland (*bracket*). Scale bars, 100 µm. **f.** Quantification of the hyperplasia of the sebaceous glands. **g, h.** Cell cycle analysis of isolated back skin keratinocytes, representative histograms (**g**) and quantification (**h**). Open bars are mean ± SD of individual samples at 1 wk post-ablation. (**a, b, f**) **p*<0.05, ***p*<0.01, ****p*<0.001, *NS*, not significant.

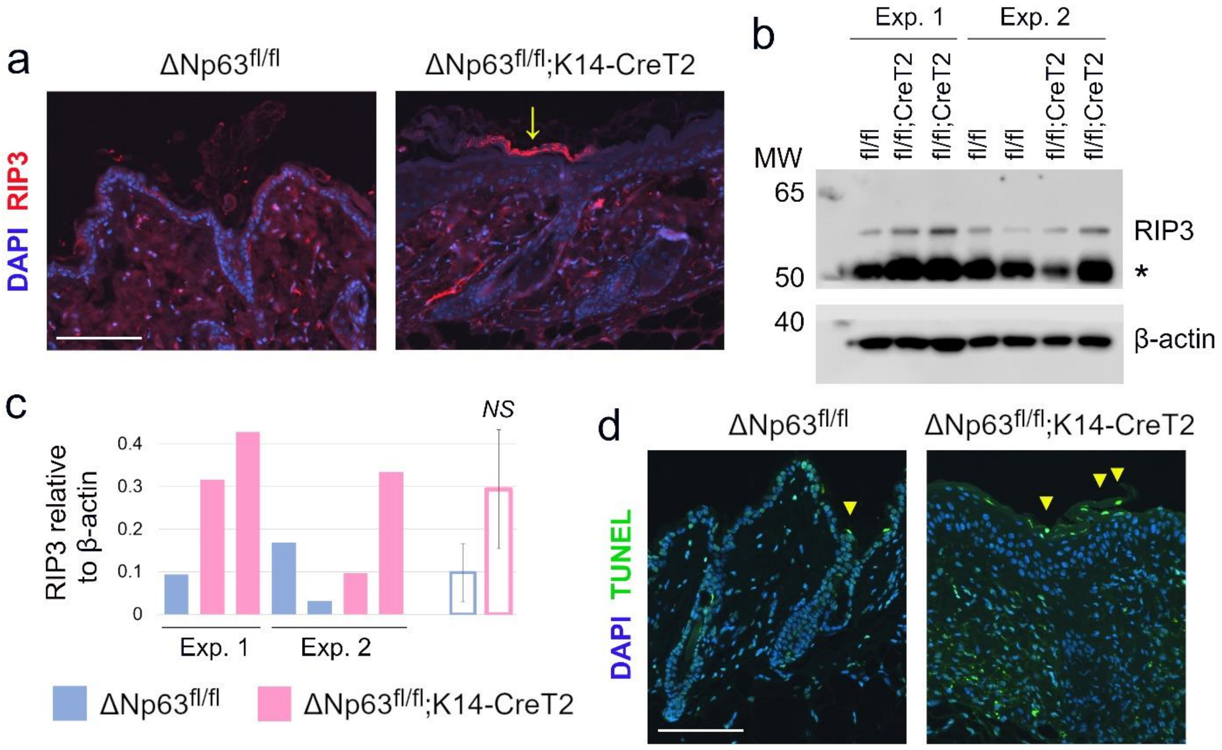


**Supplementary Figure 2**. Analysis of cell death in ΔNp63-ablated skin. **a-c.** Necrosis marker RIP3 at 1 wk post-ablation by immunofluorescent staining (**a**) and Western blot (**b**) and Western quantification (**c**). *Arrow*, RIP3 upregulation in the outermost epidermal layer (**a**). *, non-specific band (**b**). **d.** The TUNEL assay at 1 wk post-ablation. *Arrowheads*, apoptotic cells in the outermost epidermal layer. (**a, d**) Scale bars, 100 μm.

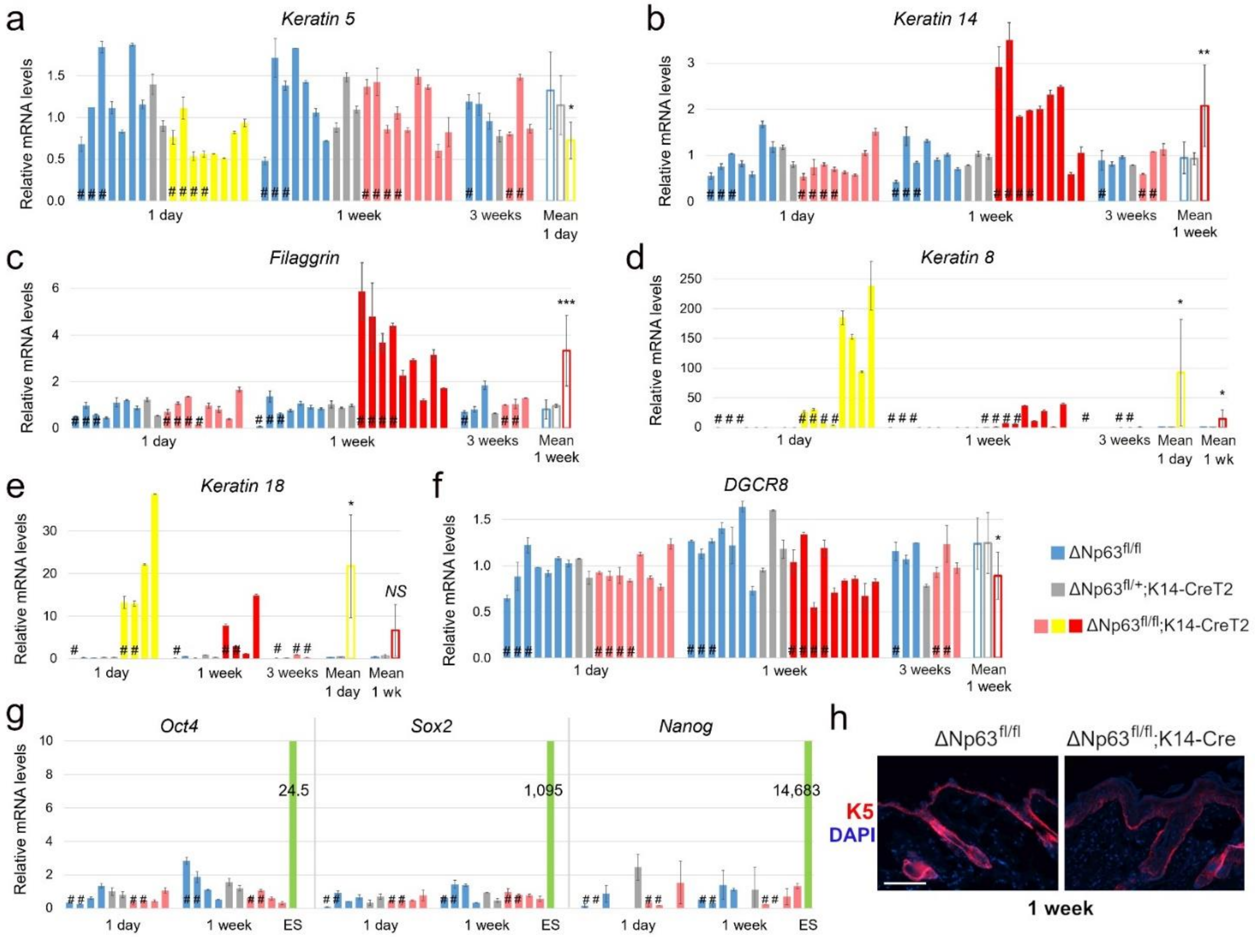


**Supplementary Figure 3**. Detailed gene expression analysis in ΔNp63-ablated skin. **a-g.** qRT-PCR of stratification markers (**a-c**), immature epidermal genes (**d, e**), *DGCR8* (**f**), and iPSC markers (**g**) on isolated primary KCs, except for # (whole back skin). Mouse ES cells served as a positive control (**g**). Mean ± SD of two technical replicas normalized to *HPRT*. Open bars are mean ± SD of individual samples at the indicated time-points. **p*<0.05, ***p*<0.01, ****p*<0.001, *NS*, not significant. **h.** Immunofluorescent staining of *Krt5*, scale bar, 100 µm.

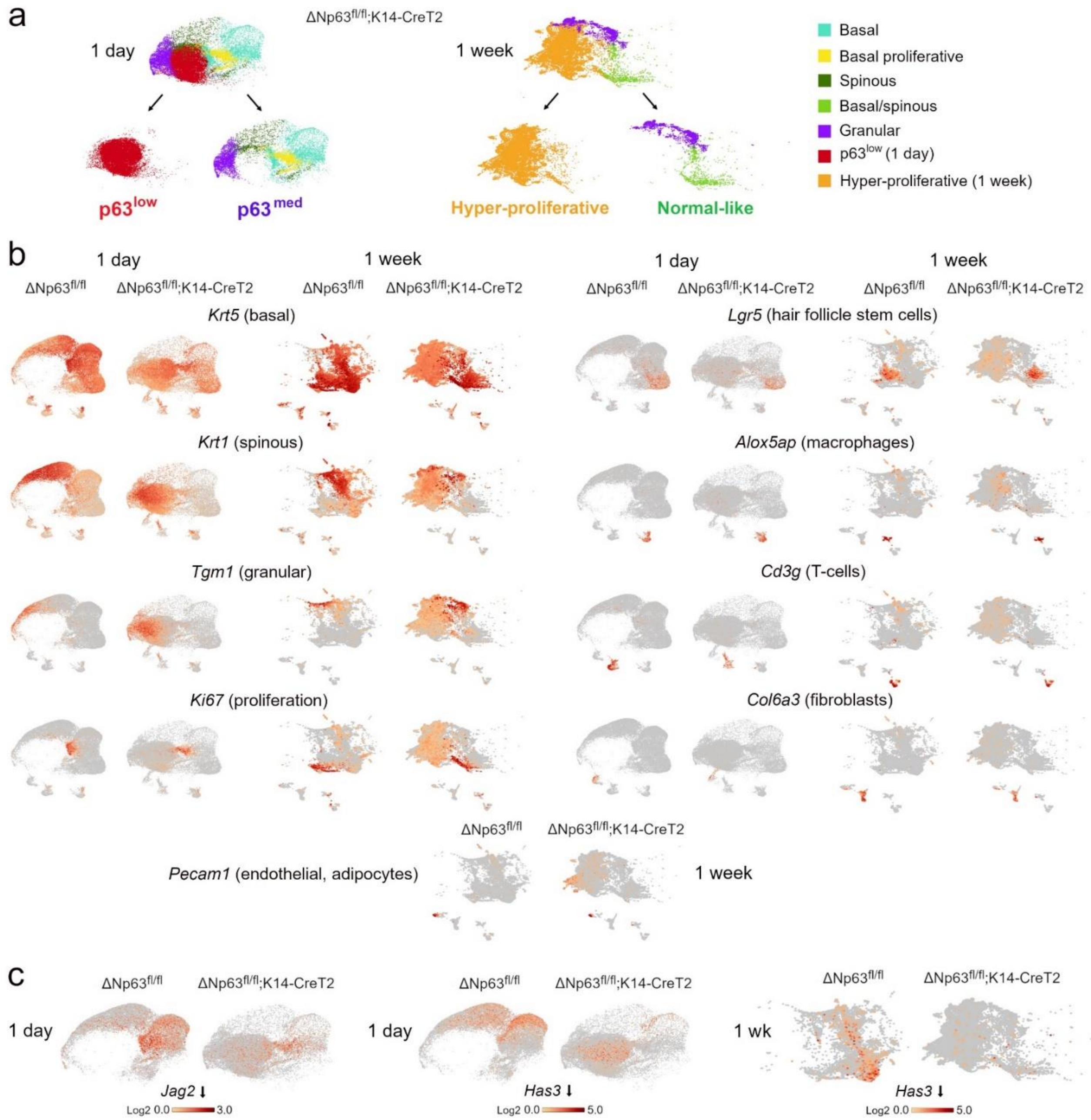


**Supplementary Figure 4**. Details of scRNA-seq analysis. **a.** Split of the UMAP plots of ΔNp63-ablated KCs at 1 day (*left*) and 1 wk (*right*) post-ablation. **b.** UMAP plots of the representative markers that were used to assign the identity of the clusters. **c.** UMAP plots of select ΔNp63 target genes.

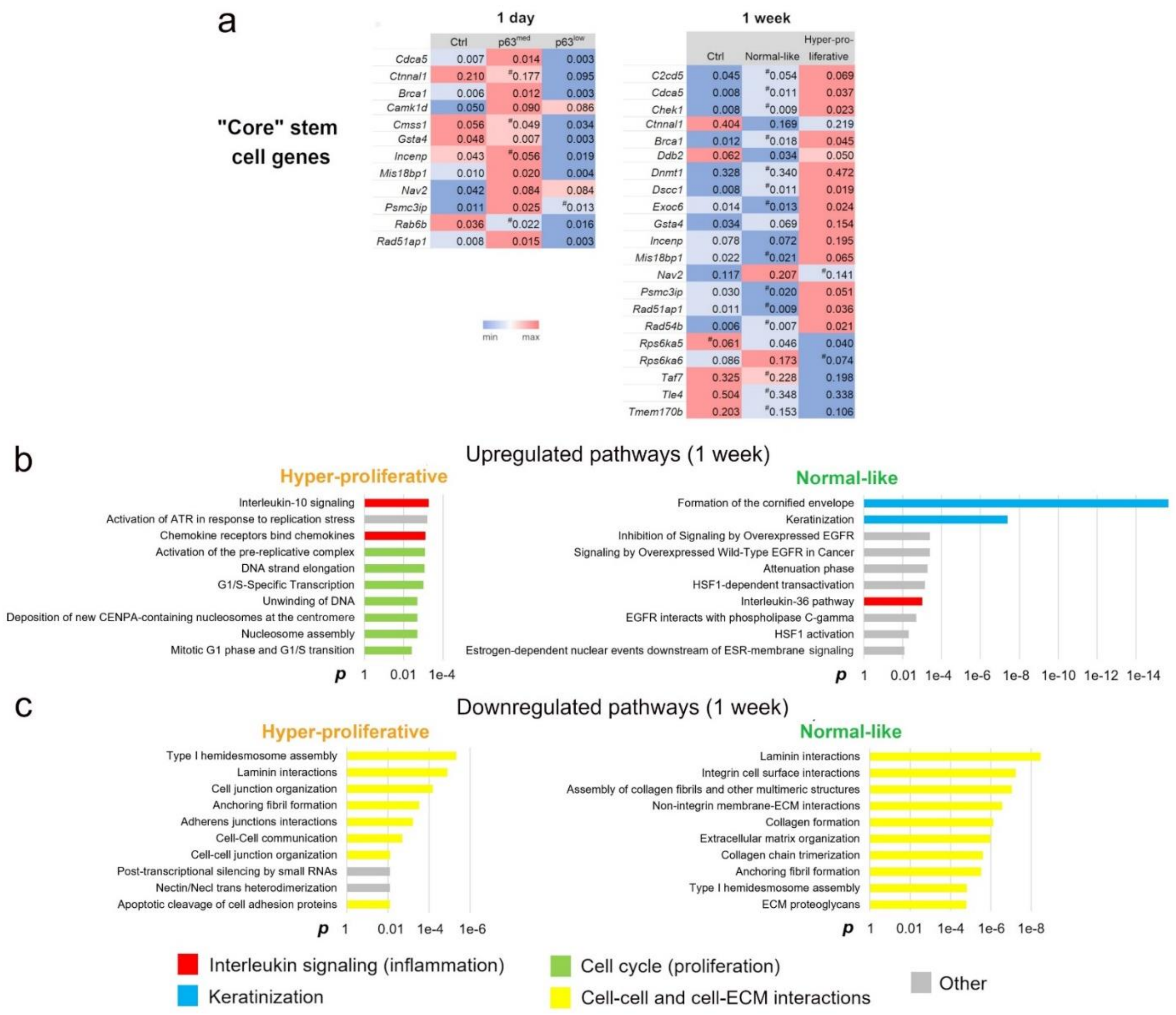


**Supplementary Figure 5**. Additional differentially regulated genes and pathways. **a.** Heatmaps of previously identified mouse "core" stem cell genes; #, not significant. **b, c.** The top ten upregulated (**b**) and down-regulated (**c**) pathways at 1 wk post-ablation, Reactome.

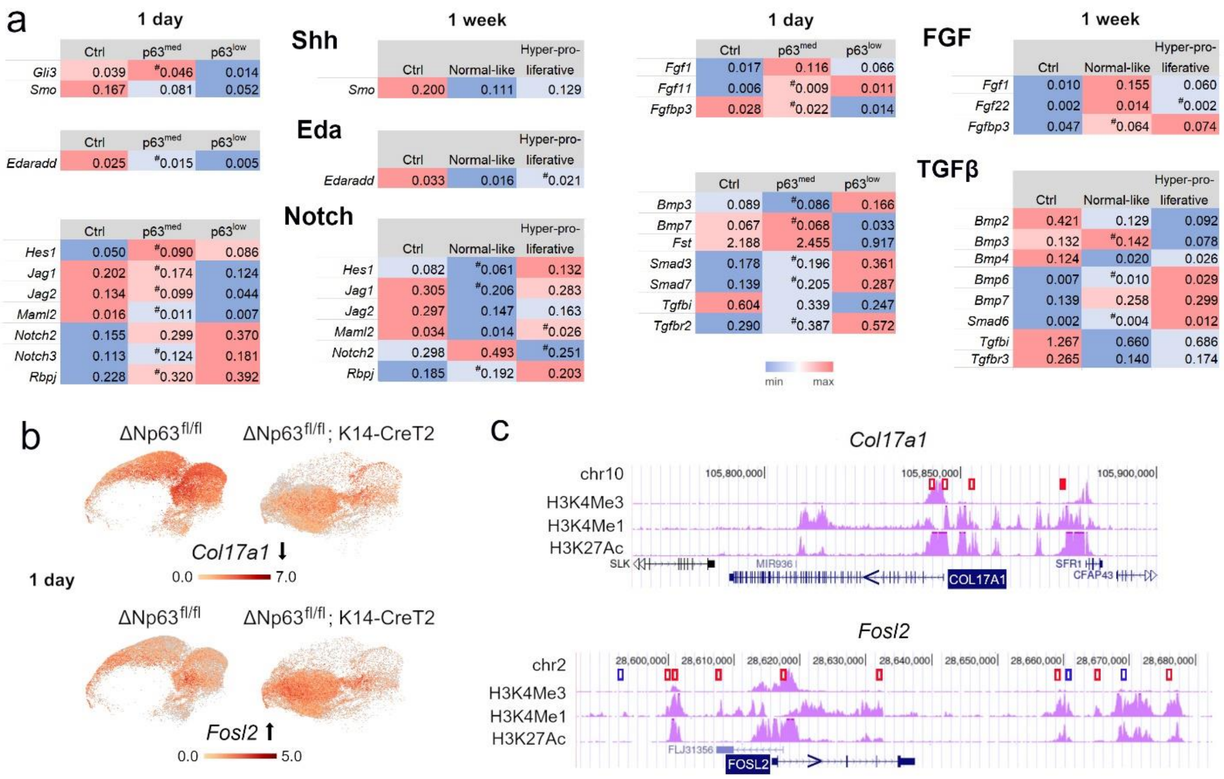


**Supplementary Figure 6**. Analysis of additional cell signaling pathways and ΔNp63 effectors. **a.** Heatmaps of the indicated signaling pathways; #, not significant. **b.** UMAP plots of *Col17a1* and *Fosl2*. **c.** Potential p63 binding sites in *Col17a1* and *Fosl2* genes from a published p63 ChIP-seq database overlayed with chromatin modification tracks (*purple*) in the UCSC genome browser, both in NHEK cells. *Red*, in all conditions; *blue*, only in proliferative conditions; *solid*, sites confirmed by ChIP-qPCR; not to scale.

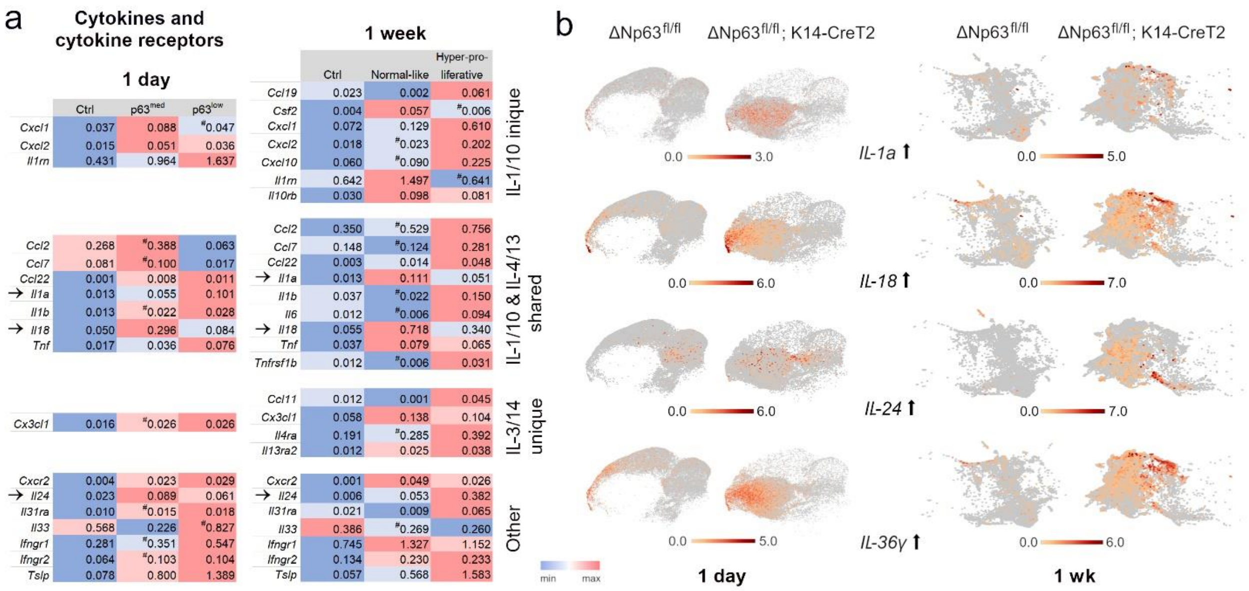


**Supplementary Figure 7**. Differentially expressed inflammatory cytokines. **a.** The heatmaps of differentially expressed genes in the indicated pathways. #, not significant. **b.** UMAP plots of the genes marked with arrows in (**a**) and *IL-36γ*.

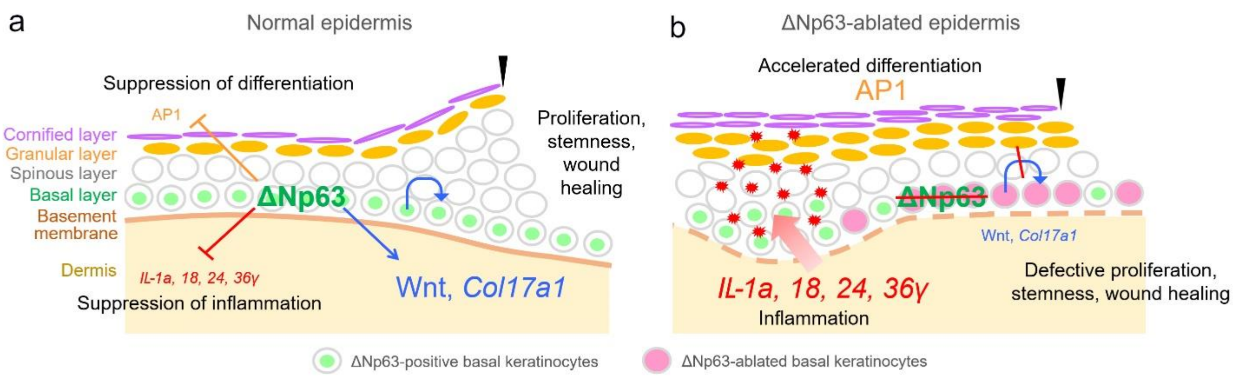


**Supplementary Figure 8**. The identified roles of ΔNp63 the adult epidermis. **a.** Normal epidermis. **b.** ΔNp63-ablated epidermis. ΔNp63 promotes KC proliferation, stemness and - as a result - wound repair via the Wnt pathway and Col17a1; restricts terminal differentiation via repression of the AP1 factors (collectively, “homeostasis”); suppresses epidermis-specific inflammatory program by repressing inflammatory cytokines. *Arrowhead*, wound border. The dashed basement membrane reflects defective cell-ECM interaction genes, not studied in detail here.

### SUPPLEMENTARY MATERIALS AND METHODS

#### Isolation of Primary Mouse Keratinocytes

Primary mouse KCs were isolated as previously described (Jensen et al. 2010) with slight modifications. Mice were euthanized by $CO_2$ asphyxiation and the back hair removed using hair clippers and Nair hair remover lotion. The skin was washed with 70% ethanol, dissected, and placed the dermal side up on a 10 cm plate on ice, the muscle and adipose tissue was carefully removed with a cell scraper and forceps. The skin was then floated epidermal side up in 10 ml 0.25% trypsin without EDTA (Gibco) either overnight at 4°C or for two hours at 37°C, transferred to a new 10 cm plate, and the epidermal layer was carefully scraped off the dermis using a cell scraper. The epidermis was then minced with a razor blade and clumps were disaggregated by pipetting in trypsin. Ice-cold PBS/10% FBS was added to the cell suspension and passed through a 40 μm mesh. Isolated KCs were gently pelleted and washed twice with ice-cold PBS/10% FBS. Cell number and viability was determined using Trypan blue (Sigma, Louis, MO) and the Beckman Coulter Z1 Counter (Beckman Coulter, Brea, CA).

#### Cell Cycle Analysis

$10^6$ primary KCs were gently pelleted, washed with ice-cold PBS, fixed in 66% ice-cold ethanol, and stored at 4°C until processing. For analysis, they underwent two rounds of washing and gentle centrifugation, were resuspended in FxCycle PI/RNase Staining Solution (Invitrogen, Waltham, MA) for 15 min. according to the manufacturer's instructions, and were analyzed with a FACSCalibur flow cytometer (BD Biosciences, Franklin Lakes, NJ) and Cyflogic software v1.2.1 (CyFlo Ltd, Turku, Finland).

**Immunohistochemistry, Immunofluorescent Staining, TUNEL Assay**

Mice were euthanized and hair was removed using hair clippers and Nair hair remover. Back skin was dissected, fixed in 10% neutral buffered formalin, embedded in paraffin, and sectioned (5 μm). Slides were deparaffinized, rehydrated, and processed for immunohistochemistry (IHC) or immunofluorescent staining. For IHC, slides were boiled in citrate antigen retrieval buffer (Vector Labs, Newark, CA) for 15 min, blocked with 5% NGS (Life Technologies, Carlsbad, CA), and incubated overnight at 4°C with primary antibodies: p63 (1:750, Cell Signaling Technology, Danvers, MA, Cat. No.13109), Ki67 (1:400, Cell Signaling Technology, Cat. No. 12202). Slides were washed in PBS and incubated with biotinylated secondary antibody (Invitrogen), followed by Vectastain Elite ABC peroxidase reagent (Vector Labs) and DAB Quanto substrate (Thermo Scientific, Waltham, MA) with hematoxylin counterstain. Additional IHC was done at Histowiz (Brooklyn, NY) with the following antibodies: BrdU (Abcam, Cambridge, UK, Cat. No. ab6326), Ki67 (Abcam, Cat. No. ab15580), p63 (GeneTex, Irvine, CA, Cat. No. GTX102425).

Immunofluorescent staining was performed as previously described (Eyermann et al. 2021) with slight modifications. Specifically, antigens were retrieved in sub-boiling buffer (10 mM Tris + 1mM EDTA + 0.05% Tween 20, pH8.0) in a microwave for 20 minutes, then incubated in 5% NGS (Life Technologies) overnight at 4°C with the following primary antibodies: Krt14 (1:800, Biolegend, San Diego, CA), Krt10 (1:200, Biolegend), Lor (1:800, Biolegend), Krt8 (1:300, Millipore, Burlington, MA), Col17a1 (1:250, Abcam, Cat. No. 186415), IL-36 (1:50, Abcam, Cat. No. ab269274), RIP3 (1:100, Santa Cruz Biotechnology, Dallas, TX, Cat. No. sc-374639). After washing in PBS, slides were incubated with fluorescent secondary antibodies. Coverslips were mounted with Prolong Gold with DAPI counterstain (Invitrogen).

The TUNEL assay was done using DeadEnd Fluorometric TUNEL System (Promega, Madison, WI, Cat. No. G3250) according to the manufacturer's instructions. Images were taken with Nikon Eclipse Ti-S microscope (Nikon, Tokyo, Japan) using NIS-Elements AR software (Nikon).

**Western Blot Analysis and Quantification**

Western blots were performed as previously described (Nemajerova et al. 2012) using the following primary antibodies: RIP3 (1:500, Santa Cruz Biotechnology, Cat. No. sc-374639), β-actin (1:1000, Santa Cruz Biotechnology, Cat. No. sc-47778). Western blot signals were quantified using the NIH Image software.

**qRT-PCR Primers**

*DGCR8*, Fwd. tacatcgactgtgcacaagg, Rev. gtatgccaagaagaacaggc ($T_m$, 82$^o$C)

*Flg*, Fwd. ggctccggatactactatg, Rev. gttcgtgctcatgctggtg ($T_m$, 88.5$^o$C)

*Krt5*, Fwd. gatccaggttctgctttatg, Rev. accctcaacaacaagtttgc ($T_m$, 83.5$^o$C)

*Krt8*, Fwd. tgttgtccatgttgctcctc, Rev. tgaacaacaagttcgcctcc ($T_m$, 83$^o$C)

*Krt14*, Fwd. caatctgcatctccaggtc, Rev. ctcctgcagatcgacaatg ($T_m$, 87$^o$C)

*Krt18*, Fwd. agaaatcgaggcactcaagg, Rev. atgatcttgctgaggtcctg ($T_m$, 80.5$^o$C)

**ChIP Primers**

*Fos* (#2), Fwd. gggtgcaagccgcctgc, Rev. taggtgggaagggcatttc ($T_m$, 85$^o$C)

*Fos* (#4), Fwd. gaggtgggattgttgacttc, Rev. gggttcatatgcaaccacag ($T_m$, 84$^o$C)

*FosB*, Fwd. gccatggtttagcaatctca, Rev. gtgctgaatcctggctgtg ($T_m$, 83.5$^o$C)

*Fzd6*, Fwd. aagaaaccacttgggaggag, Rev. caatgaggcagtcagacaac ($T_m$, 78$^o$C)

*Fzd10* (#1), Fwd. tcagtgagccaagattgtgc, Rev. gacagggtctcactttgtcc ($T_m$, 78$^o$C)

*Fzd10* (#2), Fwd. tgaactgaaatgatggcggg, Rev. attcagtcgtttgtgctgcc ($T_m$, 84.5$^o$C)

*IL-1a*, Fwd. gaactgtccttctttccctc, Rev. ctggaagttgaaaaacaagctc ($T_m$, 79$^o$C)

*IL-18*, Fwd. ttttgagaaagtctcgctctg, Rev. aatctgccacacctttggac ($T_m$, 81.5$^o$C)

*IL-24* (#3), Fwd. tctccttgaccttccttctg, Rev. aagaagaccaggtgagagtg ($T_m$, 84$^o$C)

*IL-24* (#4), Fwd. cgcaaggttaagccattctc, Rev. ctagattctaccatgtgccc ($T_m$, 81$^o$C)

*IL-36γ* (#1), Fwd. ttcccgccaattttcaaggc, Rev. tgacgtcactagctgagtac ($T_m$, 78.5$^o$C)

*IL-36γ* (#2), Fwd. gaagaatatggtcctgaggg, Rev. agctctgctcatgagaactc ($T_m$, 83.5$^o$C)

*JunB* (#1), Fwd. gctagaaacataggggcaag, Rev. gggactctctgccatactg ($T_m$, 83$^o$C)

*JunB* (#2), Fwd. aaccacagaggtgggagag, Rev. tttggtcttggcacgaggg ($T_m$, 89$^o$C)

*Wnt10a*, Fwd. gccagtgaggaataacaacc, Rev. agaccagaatccagacagtg ($T_m$, 88$^o$C)
